# Focus-Tunable Two-Photon Fiberscope Enabling *in vivo* Imaging at Selected Depths

**DOI:** 10.1101/2025.02.26.640465

**Authors:** Yuehan Liu, Jiayun L. Huang, Xingde Li

**Author notes:** These authors contributed equally to this work (Yuehan Liu, Jiayun L. Huang).

## Abstract

Miniaturized two-photon imaging devices enable real-time *in vivo* and *in situ* imaging at subcellular resolution, highly valuable for clinical applications and basic research (such as neuroscience). However, achieving high-quality volumetric imaging at varying depths remains challenging. In this study, we demonstrated a 2P fiberscope capable of three-dimensional imaging over a cylindrical volume of a 350 μm diameter and a 400 μm depth. Depth scanning was achieved by incorporating a miniature electrowetting-based varioptic lens (VL) into a two-dimensional scanning 2P fiberscope, whose focus was tuned by modulating the VL drive voltage. The performance of the fiberscope was first characterized using phantoms and then demonstrated by *ex vivo* imaging of fluorescently stained convallaria and GFP mouse brain sections, as well as *in vivo* dynamic GCaMP-based calcium imaging of cortical neurons in an awake mouse.

---

Two-photon (2P) microscopy has served as a powerful tool to image biological structures thanks to its high resolution, depth-sectioning capability, resistance to tissue scattering, and the ability to achieve deeper penetration [1–3]. Depth scan in 2P imaging has also been achieved conveniently via mechanically shifting the optics in benchtop 2P systems [4] or by using a tunable lens [5,6].

Over the past decade, miniaturized 2P imaging devices have witnessed fast development to enable *in vivo* imaging and applications for both clinical [7] and neuroscience research [8–10] based on several approaches, including the use of GRIN lens to design micro-objectives, MEMS scanners, fiber-scanning technology and/or fiber bundles. Among these technologies, the use of double-clad fiber (DCF) and tubular piezoelectric actuator for two-dimensional (2D) scanning has enabled 2P fiberscopes to fully integrate excitation and collection, achieving ultra-compact size and ultralight weight [11–14] and making it suitable for endoscopic imaging. In these systems, high transverse resolution was achieved by using large-NA objectives [15] and/or compound fiber cantilever [16].

With the goal of three-dimensional (3D) *in vivo* imaging, depth scanning in miniaturized 2P imaging devices has been actively explored [17]. Generally, two different kinds of depth scanning have been applied in miniaturized imaging systems. One method is mechanical scanning: adjusting the separation between optical elements [18], scanning the microlens [19] or the entire miniature imaging device [20,21], which has been applied to fiberscopes using shape memory alloy (SMA) wires. However, these approaches usually result in heavy weight due to complicated mechanical structural design and suffer from unsatisfactory accuracy or limited scanning range determined by the mechanical actuator. The other method is to optically tune the focal length of the imaging optics, where the lens shape can be controlled (such as fluid-based lenses and deformable elastomeric lenses) [22,23], or the refractive index of the material can be changed (such as a liquid crystal lens) to achieve adjustable focus [24].

In this study, we opted for the second approach and realized depth scanning by incorporating an electro-wetting-based variable lens (VL) into a 2P scanning fiberscope to enable focus tuning. The fiberscope weighs 1.9g, with an outer diameter of 2.4mm at the distal end, and is capable of focus tuning over a cylindrical volume field of view (FOV) with 350 μm diameter × 400 μm depth. We first evaluated its performances using fluorescent slides and microspheres, and then demonstrated the focus tuning capability using *ex vivo* samples (Convallaria and mouse brain sections) as well as *in vivo* dynamic calcium imaging of cortical neurons in an awake mouse. The low electrical power consumption, fast response, high repeatability, motionless focus scanning, and ease of operation offer tremendous benefits for *in vivo* imaging. We expect this compact, flexible 2P fiberscope with depth scanning capability will open up promising opportunities in various clinical applications and neuroscience research.

Our imaging system is based on the resonant fiber-scanner technology for performing 2D spiral scanning by the use of a tubular PZT actuator, a compound DCF cantilever, and distal-end micro-optics, as detailed in prior studies [11,13]. Given our goal of maintaining a compact, lightweight design while ensuring high image quality across depths, we adopted the optical tuning method and integrated an electro-wetting-based VL (Corning, A-25H0-D0-33) into distal-end optics. This approach not only eliminates the need for mechanical movement of optical components or the probe itself, but also preserves collection efficiency due to electro-wetting’s insensitivity to polarization, outperforming other VLs based on liquid crystal lenses. The optical power of the lens can be tuned from −35 to +35 diopters with a fine-tuning resolution of 37 mD by a driving voltage ranging from 30V to 65V. As shown in Fig. 1a and b, the curvature of the water-oil interface is adjusted by applying the voltage between the two rings of metal: one connected to the water layer, the other situated under the oil layer separated by a layer of insulator [22]. The VL has a 2.5 mm clear aperture and anti-reflective (AR) coatings optimized in the visible range. It is connected to a driver via a flexible printed circuit (FPC) cable for focus control (see Fig. 1c). The VL also has a fast response time of < 10 ms, much shorter than the frame rate of the 2D scanning (typically 2∼3 fps).

**Fig. 1.**
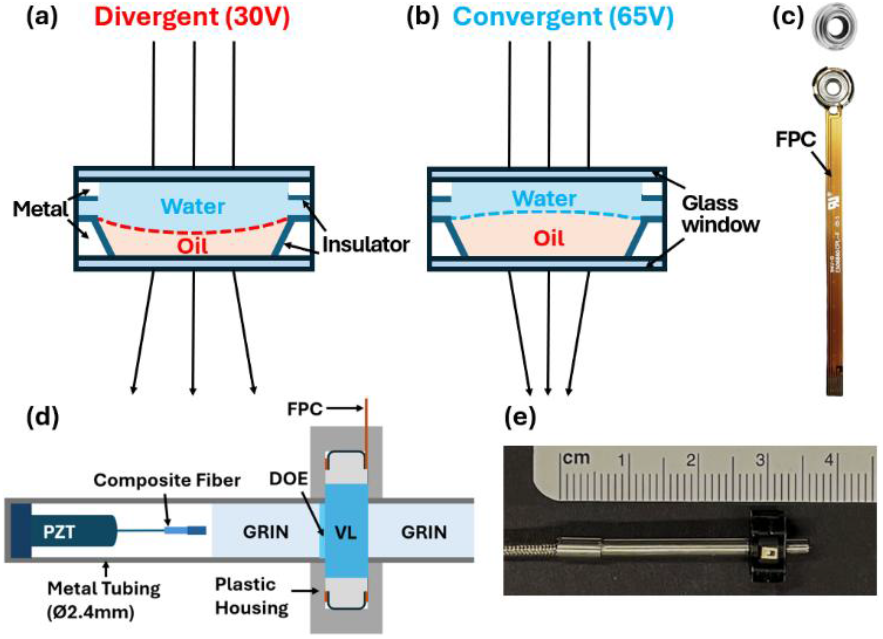
Photos and cross-sectional schematics of the electro-wetting-based VL and the focus-tuning 2P fiberscope. (a) Schematic cross-section of the A-25H0-D0-33 lens in its lowest optical power (divergent) with 30V applied between the water and the metal beyond the oil layer. (b) Schematic cross-section of the lens in its highest optical power (convergent) with 65V applied between the water and electrode. (c) Photos of A-25H0-D0-33 lens. Upper: unpackaged lens, Lower: packaged lens with FPC cable. (d) Schematic cross-section of the distal-end optics of the focus tuning 2P fiberscope. (e) Photo of the fiberscope.

The schematic of the focus-tunable 2P fiberscope is shown in Fig. 1d. The VL was positioned between two GRIN lenses (GRINTECH, LFRL-200-023-50), where the beam is nearly collimated. The performance of this configuration was confirmed by ZEMAX simulations, providing the widest range of working distances (WD). To mitigate the chromatic aberration between the 2P excitation and emission wavelengths, a diffractive optical element (DOE) was placed between the first GRIN lens and the VL, thereby enhancing collection efficiency. As shown in Fig. 1d, the distal-end optics were assembled with the composite fiber scanner within hypodermic metal tubing (2.4 mm outer diameter). Additionally, a 3D-printed plastic housing is used to protect the VL and secure its connection to the tubing assembly. The VL itself (including the ring electrode) has a diameter of 7.7 mm, while the plastic 3D-printed holder for the assembly of the VL measures 10 mm in diameter. In total, the fiberscope (including the FPC cable, metal tubing and plastic housing) weighs 1.9 g.

The WD of the fiberscope in immersion oil was measured throughout the 30-65V driving voltage range. We placed a green fluorescence reference slide on a precision motorized linear stage (Newport LTA-HS) and recorded the WD distance at each voltage. We then compared the experimentally measured WD with the simulated WD (in ZEMAX) across various voltages applied to the VL. As shown in Fig. 2, a depth scanning range of 400 μm (WD from 614 to 214 μm) was obtained. The discrepancy between the simulated and measured working distances was smallest (∼3 µm) at the lowest voltage (30V) and increased at higher voltages, reaching 24 µm at 65V. We also accounted for potential hysteresis of the VL by changing the voltage tuning direction. The VL exhibited high repeatability and no hysteresis was observed over its tunning range of -35 to +35D.

**Fig. 2.**
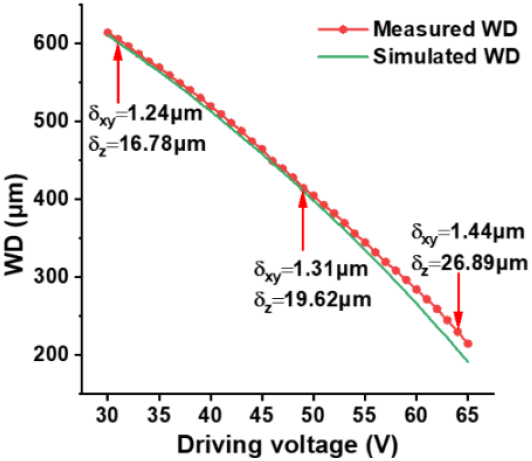
The fiberscope’s WD changing range with respect to driving voltage, and the measured resolutions at 3 different WDs.

The spatial resolution of the fiberscope was measured by imaging across 200 nm diameter fluorescent microspheres (Fluoresbrite 09834) and estimating the full width at half maximum (FWHM) of the point spread function (PSF) of an individual microsphere (Gaussian fitting for lateral resolution and Lorentzian fitting for axial resolution). Table 1 shows the measured resolution at three imaging depths in immersion oil: 230 μm (64 V), 415 μm (49 V), and 605 μm (31V). A slight reduction of resolution has been observed with increasing imaging depth. This is due to the decrease in the focusing power of the lens assembly (GRIN + VL + GRIN). Overall, the measured imaging resolution is lower than the simulation results at various depths, which might be attributed to the slight optical misalignments when assembling the fiberscope or various aberrations.

**Table 1.**
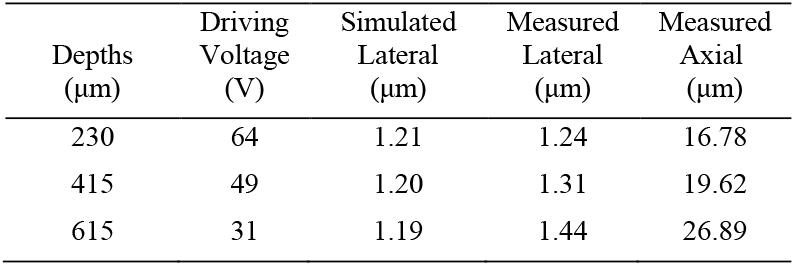
Spatial Resolution at Different Depths.

We then conducted 2P imaging with the focus-tunable fiberscope on two *ex vivo* samples: a convallaria rhizome slide (stained with acridin-orange, around 60μm thick), and a GPF mouse brain slide (around 40 μm thick). During imaging, the samples were positioned at a WD of 420 μm, corresponding to the midpoint of the depth tuning range. An FOV of a 350 μm diameter over a depth range of ∼150 μm (with a driving voltage from 42 to 55 V, in 1 V intervals for the convallaria slide and 0.5 V intervals for the brain slide) was imaged. The reconstructed volumetric image of the convallaria and mouse brain slides are presented in Fig. 3a and b, respectively, with three adjacent layers (−20, 0, +20 μm) displayed on the side.

**Fig. 3.**
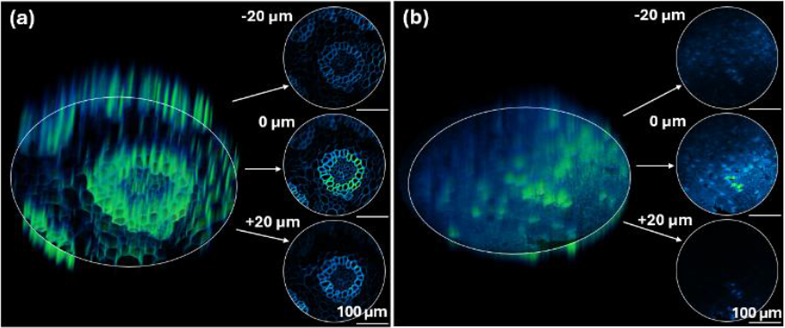
3D *ex vivo* image of Convallaria slide and mouse brain GFP slide with 2D images at the depth of the slide (denoted as 0 µm) and 20 µm above/below. (a) Convallaria slide under 10 mW excitation power at 920 nm. (b) Mouse brain GFP slide under 20 mW excitation power at 920 nm.

*In vivo* neuroactivity imaging was performed by measuring the GCaMP6m fluorescence from firing neurons through a cranial window over the somatosensory cortex on a head-fixed mouse (Jax, #005359). The overall setup is shown in Fig. 4a. During imaging, the mouse’s head was secured by using a head-restraining bar, and the 2P fiberscope was gently positioned above the cranial window with a gap of 50 μm from the #1 cover glass. All animal housing and experimental procedures were conducted according to protocols approved by the Institutional Animal Care and Use Committees (IACUC) of Johns Hopkins University.

**Fig. 4.**
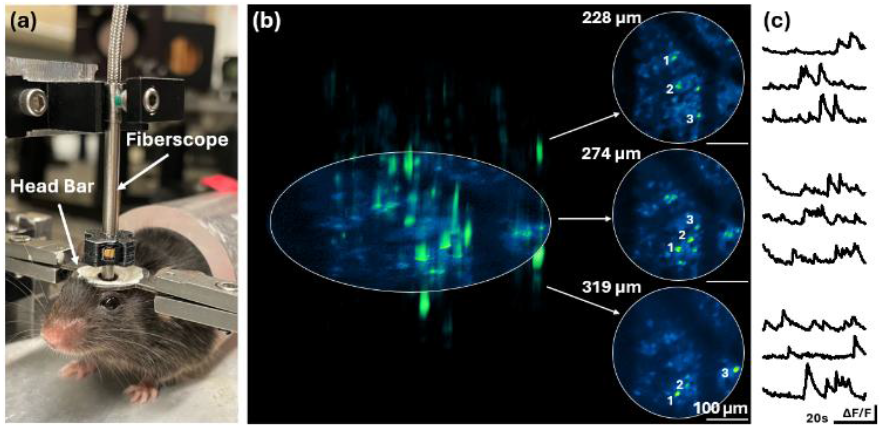
(a) Photo of the *in vivo* mouse neuroimaging set-up. (b) 3D *in vivo* calcium imaging of mouse’s cortical neurons under 35 mW excitation power at 920 nm, with 2D images at 3 different depths below the surface of the cortex. (c) ΔF/F curves of 3 representative neurons labeled at each depth.

3D imaging was performed over a 350 μm diameter FOV and a 400 μm depth (with a driving voltage from 30 to 65V, in 1 V intervals). At each depth, 100 frames of images were collected with a frame rate of 2 fps. For 3D visualization, we averaged every five frames to reduce the noise and performed maximum intensity projections to visualize all firing neurons. Fig. 4b shows the reconstructed volumetric calcium imaging of cortical neurons in an awake mouse, with three layers at depths of 228 µm, 274 µm, and 319 µm below the cortex surface displayed on the right-hand side. Individual neurons within the 3D volume are clearly resolved. The temporal calcium dynamics (ΔF/F curves) for 3 representative neurons at each layer were extracted using the CaImAn data processing pipeline [26] and illustrated in Fig. 4c.

To sum up, we demonstrated a focus-tunable 2P fiberscope by integrating an electrowetting-based VL onto a 2D resonant fiber-optic scanner, achieving a depth scanning range of 400 μm. The performance of the fiberscope was evaluated by imaging fluorescent microspheres, *ex vivo* samples and *in vivo* neurons of a mouse’s somatosensory cortex. Compared with the method with physically moving parts for depth scanning, this motionless focus scanning design is more attractive for *in vivo* and *in situ* imaging.

Based on our current design, several potential upgrades can expand its future applications. First, increasing the length of the FPC cable beyond the current commercially available 6 cm would allow the driver board to be relocated to the proximal end, facilitating easier attachment of the fiberscope to the mouse head for two-photon neural imaging in freely behaving mice. Second, leveraging the VL’s rapid response time (<10 ms), the scanning mode can be shifted from multi-layer 2D scans to rapid sequential 1D depth scans over an area, enabling time-resolved volumetric neuroimaging. Lastly, since the distal-end GRIN lens has a diameter of only 2 mm, customizing the VL packaging and holder could further reduce the overall size (and weight) of the 3D fiberscope. With advances in fabrication techniques and improved integration of the VL into scanning algorithms, we anticipate the next-generation focus-tunable 2P fiberscope to be more compact and capable of flexible volumetric imaging.

In addition to neuroscience research, this technology has the potential to ease its operation for clinical use (e.g., by eliminating the need for mechanical adjustment and shortening the detection time), and to provide substantial benefits in ensuring safe and effective surgical procedures. We hope that this focus-tunable 2P fiberscope will help unlock promising opportunities across a wide range of applications.

## Acknowledgment

The authors thank Dr. Haolin Zhang for the surgical preparation of the mouse model and Dr. Hui Lu for providing the GFP mouse brain slide.

## Data Availability Statement

Data underlying the results presented in this paper can be obtained from the authors upon reasonable request.

